# Intron retention as a novel source of cancer neoantigens

**DOI:** 10.1101/309450

**Authors:** Alicia C. Smart, Claire A. Margolis, Harold Pimentel, Meng Xiao He, Diana Miao, Dennis Adeegbe, Tim Fugmann, Kwok-Kin Wong, Eliezer M. Van Allen

**Author notes:** These authors contributed equally to this work (Alicia C. Smart, Claire A. Margolis). Correspondence to (EMV).

## Abstract

Personalized cancer vaccine strategies directed at tumor neoantigens derived from somatic mutations in the DNA are currently under prospective evaluation^1, 2^. Alterations in tumor RNA, rather than DNA, may also represent a previously-unexplored source of neoantigens. Here, we show that intron retention, a widespread feature of cancer transcriptomes^3, 4^, represents a novel source of tumor neoantigens. We developed an *in silico* approach to identify retained intron neoantigens from RNA sequencing data and applied this methodology to tumor samples from patients with melanoma treated with immune checkpoint blockade^5, 6^, discovering that the retained intron neoantigen burden in these samples augments the DNA-derived, somatic neoantigen burden. We validated the existence of retained intron derived neoantigens by implementing this technique on cancer cell lines with mass spectrometry-derived immunopeptidome data^7, 8^, revealing that retained intron neoantigens were complexed with MHC I experimentally. Unexpectedly, we observed a trend toward lack of clinical benefit from immune checkpoint blockade in high retained intron load-tumors, which harbored transcriptional signatures consistent with cell cycle dysregulation and DNA damage repair. Our results demonstrate the contribution of transcriptional dysregulation to the overall burden of tumor neoantigens, provide a foundation for augmenting personalized cancer vaccine development with a new class of tumor neoantigens, and demonstrate how global transcriptional dysregulation may impact selective response to immune checkpoint blockade.

**Statement of significance:** We developed and experimentally validated a computational pipeline to identify a novel class of tumor neoantigens derived from RNA-based intron retention, which is prevalent throughout cancer transcriptomes. The discovery of transcriptionally-derived tumor neoantigens expands the tumor immunopeptidome and contributes potential substrates for personalized cancer vaccine development.

---

Cancer immunotherapy relies on immune recognition of tumor cells as foreign, enabling an immune response that destroys tumor cells^9^. Recent clinical successes of immune checkpoint blockade therapies across tumor types demonstrate the ability of these agents to enhance immune destruction of tumors in patients with advanced disease^10, 11^. In addition, personalized cancer vaccines, alone or in combination with immune checkpoint blockade therapies, have shown potential as targeted immunotherapies^1, 2^. Tumor neoantigens, which arise from somatic nonsynonymous mutations in expressed genes, generate novel peptides that are foreign to the immune system and are able to stimulate an immune response^5, 12–14^. Genes that are not normally expressed in adult somatic tissues, such as cancer germline antigens, also have been shown to generate immunogenic peptides that contribute to tumor rejection^15, 16^.

Analysis of untreated tumor transcriptomes demonstrates abundant dysregulation of RNA splicing, characterized predominantly by intron retention, even in the absence of somatic mutations affecting the splicing machinery^3, 4^. Intron retention occurs when the spliceosome fails to remove an intron from the pre-mRNA transcript, causing it to remain in the final mRNA transcript. Products of aberrant splicing events, including intron retention, are endogenously processed, proteolytically cleaved into 8–11 amino acid peptides, and presented on the cell surface bound to MHC class I molecules for recognition by CD8 T cells^17^. Transcripts containing retained introns often enter the nonsense-mediated decay (NMD) pathway and usually do not lead to expression of full-length proteins due to premature stop codons contained in the intronic region^18^. The NMD pathway relies on recognition of these premature termination codons, which requires that transcripts undergo translation in order to be targeted for degradation. Peptides generated through this pioneer round of translation are a source of antigens presented by the MHC class I pathway, as evidenced by the finding that presentation of an antigen can occur even when synthesis of its full-length protein is disrupted^19^. In addition, genes yielding transcripts likely to undergo NMD have been found to preferentially give rise to MHC class I-associated peptides, and the likelihood of undergoing NMD is predictive of MHC class I-associated peptide generation^20^. These findings suggest that aberrant peptide products generated through the translation and degradation of retained introns may be a novel source of tumor neoantigens; however, thus far there is no direct evidence that retained introns result in tumor neoantigens presented through the MHC class I pathway, nor is there an established relationship between this form of transcriptional dysregulation and clinical benefit from immune checkpoint blockade therapies.

In this study, we develop a computational method to identify putative retained intron neoantigens in tumor samples from two clinical cohorts of melanoma patients treated with immune checkpoint blockade therapies. We then apply our method to a set of cancer cell lines with corresponding mass spectrometry-derived immunopeptidome data and show that putative retained introns identified by our pipeline are found experimentally in complex with MHC I molecules. Further, we explore trends associated with retained intron (RI) neoantigens versus somatic neoantigens and the relevance of RI neoantigens to patient clinical outcome. These findings provide evidence that RI neoantigens are detectable in tumor cells and contribute to tumor immunogenicity.

To identify putative RI neoantigens in tumor samples, we developed a computational pipeline using bulk RNA sequencing (RNA-Seq) data from clinical tumor samples as input (Fig. 1A, Methods). The pipeline leverages novel adaptations of established methods and publicly-available databases^21–26^ to create a standardized workflow optimized for identification of putative RI neoantigens. Briefly, expressed intron retention events were detected from pre-processed tumor sample RNA-Seq data, and intron fragments likely to be translated into peptides based on their position downstream from a translated exon and upstream from an in-frame stop codon were identified. Predicted binding affinities between RI peptide sequences and sample-specific HLA Class I alleles were calculated in order to identify candidate RI neoantigens. Critically, preliminary results were then filtered and thresholded to exclude artifacts generated by sources including erroneous transcriptome annotations, low sequencing coverage, and events that are unlikely to stimulate immune response due to tolerance mechanisms such as overlap with normal protein sequences and intron retention events detected in normal tissue. This process (Methods) generated a robust final list of putative RI neoantigens for each sample.

**Figure 1.**
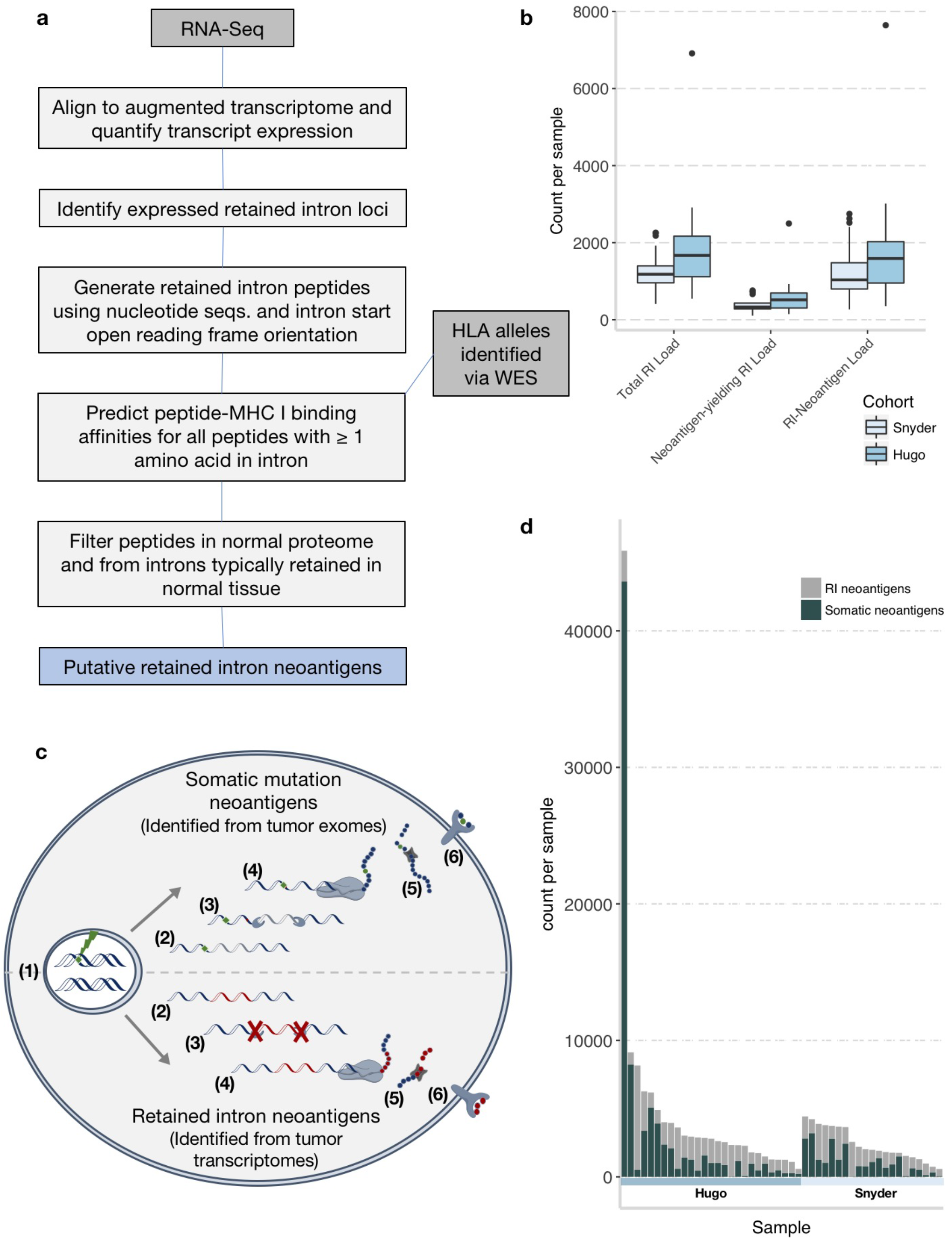
Computational identification of RI neoantigens significantly augments overall neoantigen burden in the Hugo and Snyder patient cohorts. **A**, *In silico* retained intron pipeline detects intron retention events from whole transcriptome sequencing, determines open reading frames extending into intronic sequences, and identifies putative HLA-specific neoantigens. **B**, Distribution of total RI load, neoantigen-yielding RI load, and RI neoantigen load in patient cohorts (n = 27 Hugo, n = 21 Snyder). Boxplots show the median, first and third quartiles, whiskers extend to 1.5 x the interquartile range, and outlying points are plotted individually. **C**, Neoantigen presentation pathway: Somatic DNA mutations (1) are transcribed (2), spliced (3) and missense mutations are translated (4) and undergo processing into 9-10mer peptides (5), which are presented on the cell surface through the MHC I pathway (6). RI neoantigens are produced from unmutated DNA (1), transcribed (2), and undergo defective splicing resulting in intron retention (3). RI transcripts are translated resulting in abnormal peptides and early termination (4). Abnormal proteins are degraded through the NMD pathway, processed into 9-10mer peptides (5), and presented on the cell surface through the MHC I pathway (6). **D**, Somatic and RI-neoantigen load for each patient, sorted by clinical cohort.

We applied this pipeline to tumor samples from two published cohorts of melanoma patients who received immune checkpoint blockade therapies^5, 6^ to identify putative RI neoantigens (n = 48 melanomas; Supplementary Tables S1 and S2). Another clinical immunotherapy-treated melanoma cohort^27^ was excluded from this analysis because it was generated from formalin-fixed, paraffin-embedded (FFPE) tumor samples and sequenced using a transcriptome capture technique, which did not result in adequate preservation of RI transcripts (unpublished data).

Both cohorts had comparable levels of intron retention and RI neoantigens, with one outlier in the Hugo et al. cohort with abundant transcriptional dysregulation and significantly elevated retention and RI neoantigen burden, defined henceforth as the count of unique RI-derived neoantigen peptide sequences per individual (Fig. 1B). However, slight variation in RI neoantigen load between cohorts was expected given differences in sequencing run, depth, and quality, which can be especially apparent in the context of RNA-Seq analysis^28^. Total number of retained introns was tightly correlated with RI neoantigen load in both cohorts (R^2^ = 0.93 and 0.86 for the Hugo and Snyder cohorts, respectively), with a mean of ~3 neoantigens arising from each neoantigen-yielding retained intron for both cohorts (Supplementary Fig. S1 and Supplementary Table S1).

To determine the total neoantigen load for each patient, we considered neoantigens arising from both RIs (RNA-based), generated via our pipeline, and somatic mutations (DNA-based), generated via published methods^27^ (**Fig. 1C**, Supplementary Table S1). The majority of patients from both cohorts showed significantly augmented total neoantigen loads with the incorporation of RI neoantigens into the analysis. Mean somatic neoantigen load across cohorts was 2,218 and mean RI neoantigen load was 1,515, yielding a ~0.7-fold increase in mean total neoantigen load with the addition of RI neoantigens (**Fig. 1D**). There was not a significant correlation between somatic neoantigen load and RI neoantigen load (p = 0.63) (Supplementary Fig. S2), suggesting that analysis of the complete immunopeptidome may provide greater insight into patient response to immunotherapy.

To experimentally demonstrate that RIs are endogenously processed and presented through the MHC class I pathway, we identified RI neoantigens in tumor cell lines that were found complexed to MHC I. We utilized RNA-Seq data from multiple human tumor cell lines and their corresponding MHC I mass spectrometry data elucidating their immunopeptidomes to query for *in vitro* presentation of computationally-predicted RI neoantigens^7, 8^ (Supplementary Table S3). In MeWo, a melanoma cell line, the RI neoantigens *EVYAAGKYV* and *YAAGKYVSF* were both predicted with our pipeline and experimentally discovered in complex with MHC I via mass spectrometry with high confidence (**Fig. 2A**). Both of these RI neoantigens are predicted to arise from a retained intron in the gene *KCNAB2* at genomic locus chr1:6142308-6145287. Similarly, we found RI neoantigens identified with both methods in another melanoma cell line, SK-MEL-5 (RI neoantigens *AMSDVSHPK* and *LAMSDVSHPK* from an intron in gene *SMARCD1*), in B cell lymphoma cell lines CA46 (RI neoantigen *FRYVAQAGL* from an intron in gene *LRSAM1)* and DOHH-2 (RI neoantigens *TLFLLSLPL* and *FLLSLPLPV* from an intron in gene *CYB561A3*), and in leukemia cell lines HL-60 (RI neoantigen *SVLDDVRGW* from an intron in gene *TAF1)* and THP-1 (RI neoantigen *LTSQGKSAF* from an intron in gene *ZCCHC6)* (**Fig. 2B** and Supplementary Fig. S3). Additionally, the same procedure was performed using computationally-derived, somatic mutation neoantigens, and comparable rates of mass spectrometric peptide detection were observed in this setting (Supplementary Table S4). The discovery of peptides in complex with MHC I *in vitro* via mass spectrometry with sequences shared by RI neoantigens predicted computationally with our pipeline provides direct evidence of the processing and presentation of RI neoantigens through the MHC I pathway.

**Figure 2.**
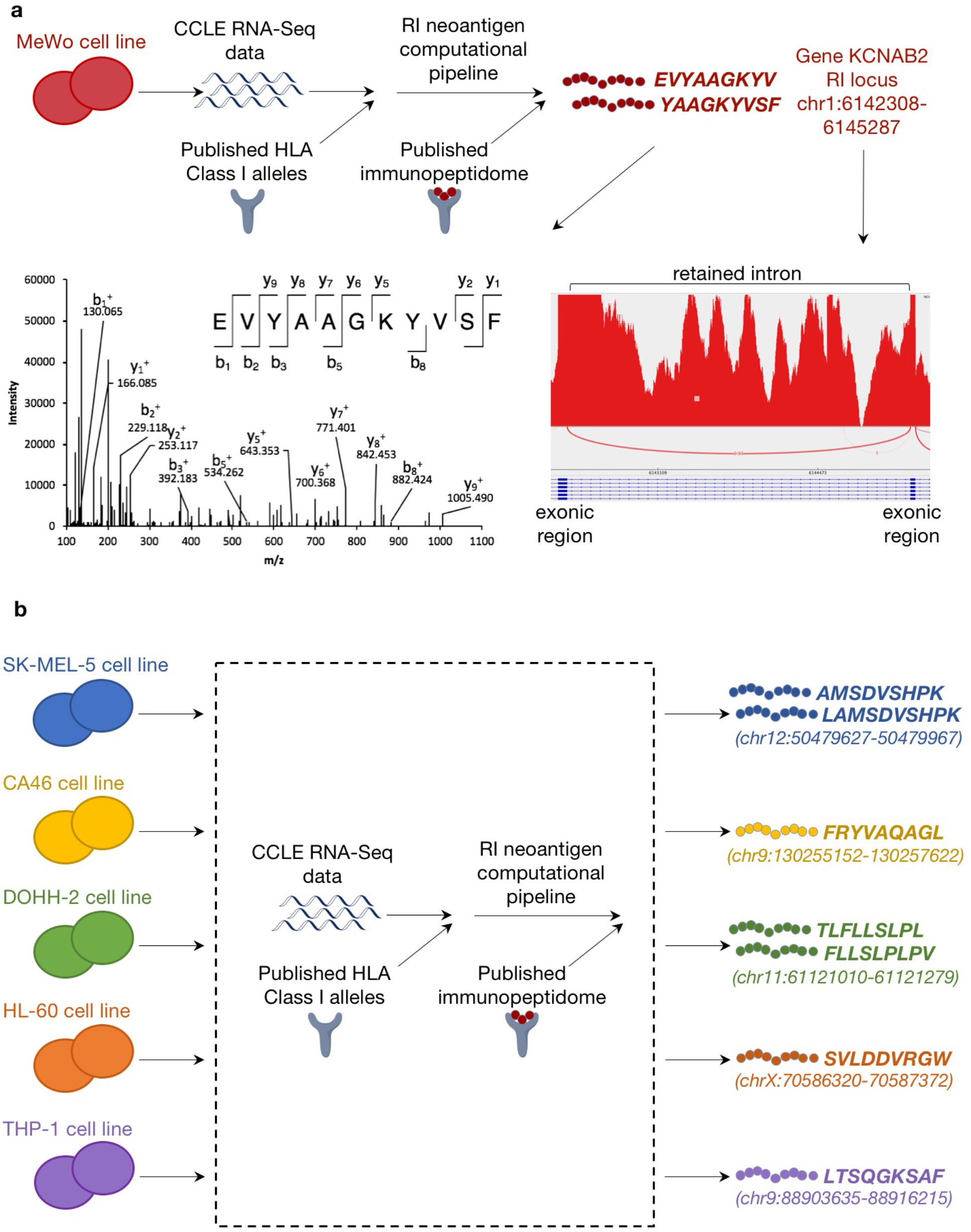
Predicted RI neoantigens in cancer cell lines are experimentally identified in complex with MHC Class I. **A**, The application of our method to the MeWo cell line revealed two RI neoantigens originating from the same intron in gene *KCNAB2* which were both predicted *in silico* and found by mass spectrometry to be present in the MeWo immunopeptidome. Corresponding Integrative Genomics Viewer (IGV) sashimi plot indicating RNA-Seq read depth in intron and surrounding exons (RI expression in TPM=5.13, percent-spliced-in [PSI] value=1.07%) as well as mass spectra for the RI neoantigen peptides, are shown. **B**, A similar procedure revealed RI neoantigens predicted by our pipeline with mass spectrometric evidence supporting their presentation on cell surfaces in complex with MHC Class I in five other human tumor cell lines: SK-MEL-5, CA46, DOHH-2, HL-60, and THP-1.

To explore the role of RI neoantigens in predicting individual response to checkpoint blockade, we next analyzed predicted RI neoantigens in melanoma patient cohorts (**Fig. 1B, D**). We hypothesized that a subset of RI neoantigens may be differentially expressed between patients who did and did not respond to immune checkpoint blockade therapy. Such neoantigens might have both therapeutic relevance for cancer vaccines and clinical relevance as biomarkers for potential benefit of immune checkpoint blockade therapy. We identified a total of 6,178 responder-exclusive RI neoantigens, or RI neoantigens that appeared in at least one responder and no nonresponders from either cohort, with 1,017 of those present in two or more responders and 398 present in at least one patient from each cohort (p > 0.05 for all, Methods) (**Fig. 3A, B** and Supplementary Table S5). The most prevalent responder-exclusive RI neoantigen, *LPVSTLPPSL*, arose from a RI within the *TRIP6* gene and was expressed in seven responder samples from the Hugo cohort with a median expression level of 9.38 transcripts per million (**Fig. 3C**). No non-responder samples from either cohort were predicted to have the *LPVSTLPPSL* RI neoantigen. Additionally, another RI in the *TRIP6* gene yielded a second responder-exclusive RI neoantigen, *RPDRQVTLPL*, which was expressed in four samples from the Hugo cohort with a median expression level of 11.01 transcripts per million. The implications of expression of these RI neoantigens as well as other responder-exclusive RI neoantigens in patient samples merit further study in larger clinically-annotated cohorts.

**Figure 3.**
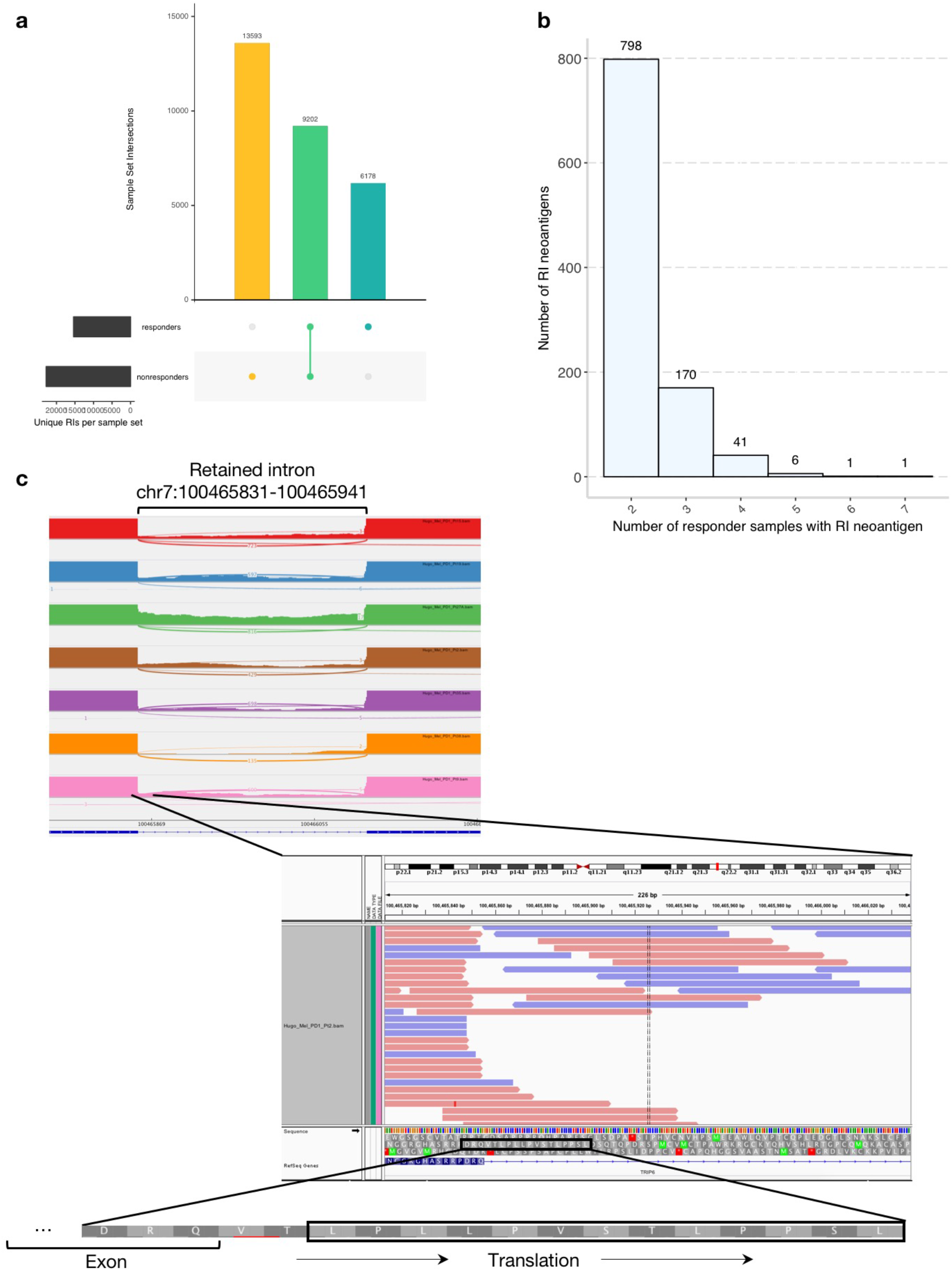
A subset of RI-neoantigens were exclusive to patients who received clinical benefit from immune checkpoint blockade therapy (n = 22 patients total). **A**, UpSet visualization of set intersections for RI neoantigens exclusive to responders, nonresponders, and shared by at least one patient in both response groups. **B**, Distribution of responder-exclusive RI neoantigens across patients. Each bar corresponds to the number of unique RI neoantigens common to the indicated number of responder patients. The most common responder-exclusive RI neoantigen was present in seven patients. **C**, IGV sashimi plot highlights RNA-Seq read depth for the seven patients with responder-exclusive RI neoantigen *LPVSTLPPSL*. Expression in TPM and PSI (percent spliced-in) values for each patient as follows, from top to bottom of stacked sashimi plots: Pt15 TPM=4.66, PSI=4.10%; Pt19 TPM=10.96, PSI=13.90%; Pt27A TPM=9.87, PSI=8.90%; Pt2 TPM=12.93, PSI=18.05%; Pt35 TPM=5.80, PSI=3.95%; Pt38 TPM=1.09, PSI=3.91%; Pt9 TPM=9.38, PSI=7.88%. The zoomed-in region of a representative sample expressing the retained intron shows the identified RI-neoantigen amino acid sequence, translated in the correct reading frame from the previous exon. An N-terminal portion of the retained intron is likely translated due to its position upstream of an in-frame stop codon (not shown).

Given that somatic neoantigen burden is a known correlate of clinical benefit from immune checkpoint inhibitor therapy in melanoma^27^, we then examined whether RI neoantigen load might be similarly associated in our cohorts of melanoma patients. However, there was no significant association between RI neoantigen load and clinical benefit from immune checkpoint blockade therapy in either cohort alone or in aggregate, nor was there correlation with expression of canonical markers of immune cytolytic activity, CD8A, GZMA, or PRF1^29^ (p > 0.05 for all, **Fig. 4A** and Supplementary Fig. S4). Further, patients who did not respond to immune checkpoint blockade therapy tended to have higher RI neoantigen loads than those who did respond, although this trend was not statistically significant (p = 0.29 and 0.61 for the Snyder and Hugo cohorts, respectively). There was no association between other clinical covariates (i.e., age, sex, disease status) and RI neoantigen load (Supplementary Fig. S5).

**Figure 4.**
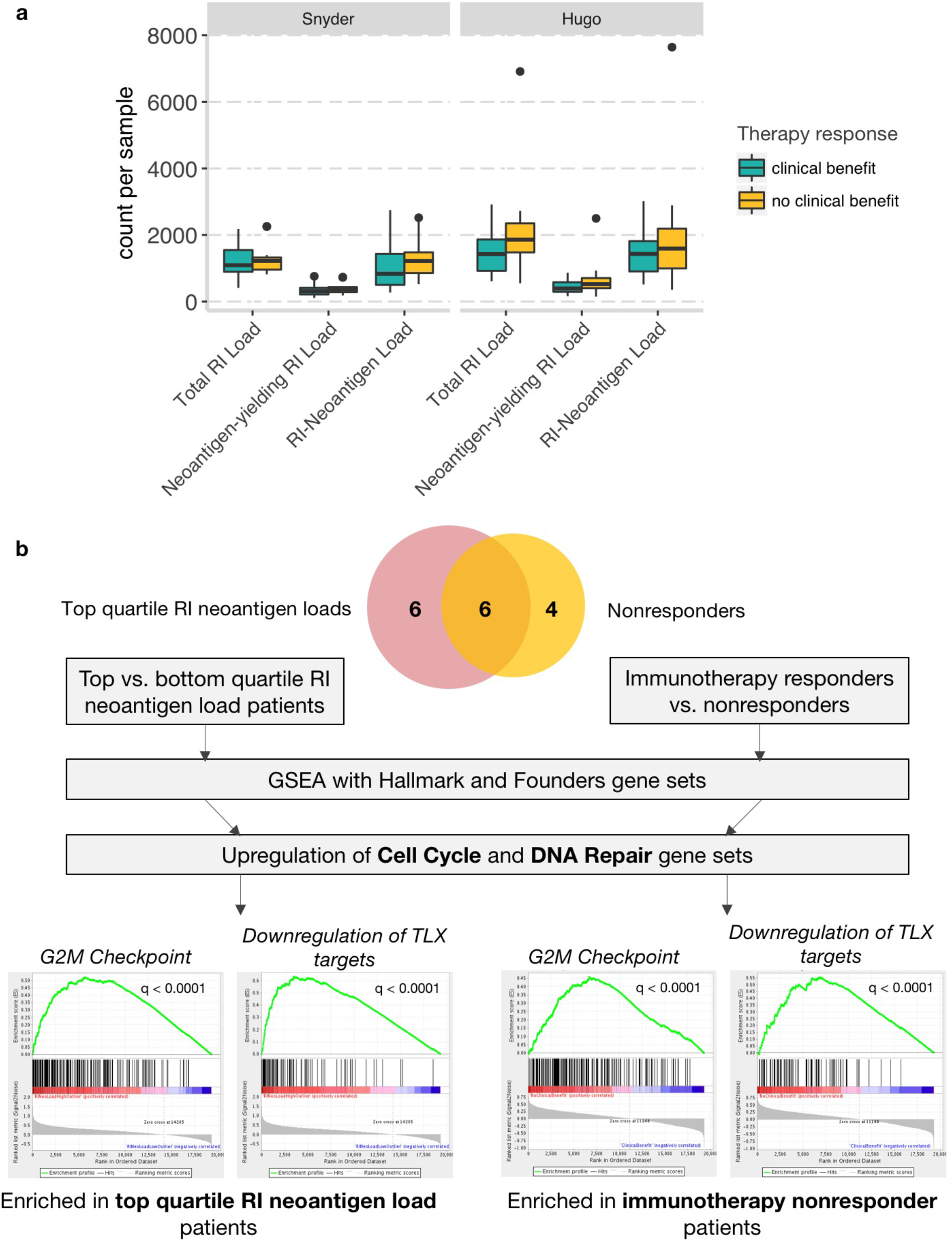
Patients with high RI neoantigen loads and immunotherapy nonresponders show enrichment of similar transcriptional programs. **A**, Association of RI load, neoantigen-yielding RI load, and RI neoantigen load with clinical benefit from immunotherapy in Hugo (n = 14 clinical benefit, n = 13 no clinical benefit) and Snyder (n = 8 clinical benefit, n = 13 no clinical benefit) patient cohorts. Both cohorts show a nonsignificant trend (p > 0.05, two-sided Mann-Whitney U test) towards association between RI neoantigen load and immunotherapy nonresponse. Boxplots show the median, first and third quartiles, whiskers extend to 1.5 x the interquartile range, and outlying points are plotted individually. **B**, Gene Set Enrichment Analysis (GSEA) was performed comparing top vs. bottom quartile RI neoantigen load patients and immunotherapy nonresponders vs. responders. Only half of the top quartile RI neoantigen load patients were overlapping as nonresponders to immunotherapy. Enrichment of cell cycle- and DNA repair-related gene sets was seen in both high RI neoantigen load patients and immunotherapy nonresponders. Representative GSEA enrichment plots from the *G2M checkpoint* and *Downregulation of TLX targets* gene sets are shown for both the top vs. bottom quartile RI neoantigen load patients and immunotherapy nonresponders vs. responders comparisons. FDR q-values are indicated on plots.

To investigate this paradoxical trend and the transcriptional correlates of RI neoantigen burden, we performed gene set enrichment analysis (GSEA)^30^ on the top (n = 12) vs. bottom (n = 11) quartile patients by RI neoantigen load across both patient cohorts. Analysis of 50 “Hallmark” gene sets representing major biological processes^30^ revealed a statistically significant enrichment in expression of cell cycle-related genes, including those linked to the G2M checkpoint (q < 0.0001), E2F targets, which play a role in the G1/S transition of the cell cycle (q < 0.0001), MYC targets (q = 0.012, q = 0.026), and mitotic spindle (q = 0.016), in the top quartile RI neoantigen load patients compared to the bottom quartile patients (**Fig. 4B** and Supplementary Table S6). In further GSEA analysis of the more refined “Founders” gene sets describing the significantly enriched Hallmark gene sets, the most strongly enriched gene sets in the top quartile RI neoantigen load patients were related to DNA replication and damage repair, e.g., downregulation of *TLX* targets including tumor suppressor genes *CDKN1A, SIRT1*, and *PTEN* (q < 0.0001)^31^, activation of ATR in response to replication stress (q < 0.0001), DNA dependent DNA replication (q < 0.0001), *BRCA* centered network (q < 0.0001) (**Fig. 4B** and Supplementary Table S6).

Interestingly, similar results were seen when performing GSEA on the same set of patients but grouped based on response to immunotherapy (n = 10 nonresponders and n = 13 responders), despite the fact only half of the top quartile RI neoantigen load patients were nonresponders (**Fig. 4B**). Compared to responders, immunotherapy nonresponders showed statistically significant enrichment in many of the same gene sets related to cell cycle and DNA damage repair as were upregulated in the top quartile RI neoantigen load patients for both the Hallmark and Founders gene sets: G2M checkpoint (q < 0.0001), E2F targets (q < 0.0001), mitotic spindle (q < 0.0001), MYC targets (q = 0.005, q = 0.104), downregulation of *TLX* targets (q < 0.0001), activation of ATR in response to replication stress (q = 0.001) (**Fig 4B** and Supplementary Table S6). Our results reveal a transcriptional similarity between high RI neoantigen load patients and immunotherapy nonresponders, and suggest that cell cycle dysregulation may be influencing both global aberrant RNA splicing and immunotherapy response, potentially via distinct biological mechanisms and pathways.

This study establishes a novel and previously uninvestigated source of neoantigens derived from RNA-based tumor events. We demonstrate that tumor-specific RI neoantigens can be identified computationally in both patient- and cell line-derived samples and a subset can be validated as presented on the cell surface in complex with MHC I. We developed a computational framework to identify patient-specific neoantigens arising from intron retention events and identified RI neoantigens in tumor samples from two clinical cohorts of melanoma patients. Putative RI-neoantigen peptides predicted *in silico* from multiple human tumor cell lines were found experimentally to be bound to the MHC Class I molecule *in vitro* through mass spectrometry. These data support the hypothesis that aberrant splicing results in intron retention, which generates abnormal transcripts that are translated into immunogenic peptides and presented to the immune system, underscoring their relevance in patients receiving immunotherapy. Notably, further studies and experimental approaches will be necessary to clinically validate the immunogenicity of specific RI neoantigens in patients, including identification of T cells specific to predicted RI neoepitopes.

Additionally, although RI neoantigen load is not predictive of response to immune checkpoint blockade therapy as a global measure, a subset of RI neoantigens are associated with treatment response and may have further clinical relevance for both cancer vaccine formulation and immunotherapy response prediction. Moreover, we discovered that patients with top quartile RI neoantigen loads are transcriptionally similar to immunotherapy nonresponders; both patient groups have enrichment of cell cycle and DNA damage repair-related gene sets. Intron retention has been shown to regulate the cell cycle in both non-malignant^32^ and malignant cells^33^. Further, high RI neoantigen load and nonresponse to checkpoint blockade were associated with downregulation of *TLX* target genes including tumor suppressors *CDKN1A, PTEN*, and *SIRT1*, genes that are involved in halting cell cycle progression in response to DNA damage sensing^34^. These findings are provocative given the emerging synergistic relationship between cell cycle inhibition and immune checkpoint blockade therapies^35–37^. Small molecule cell cycle inhibitors, which reduce activity of *E2F* targets, have been shown to enhance tumor cell antigen presentation as well as inhibit proliferation of immunosuppressive regulatory T cells, possibly explaining their potentiation of the effects of checkpoint blockade. In our cohorts, both high RI neoantigen load and checkpoint inhibitor nonresponse were associated with increased expression of *E2F* targets. Further investigation may elucidate whether a similar mechanism underlies the paradoxical trend between elevated RI neoantigen burden and immunotherapy nonresponse.

Identification of a wider array of tumor neoantigens, including those derived from somatic mutation, aberrant gene expression, and splicing dysregulation, will contribute to a more complete understanding of the tumor immune landscape. Additional work dissecting the relationship between the prediction, processing and presentation, and ultimate immunogenicity of neoantigens derived from different sources will be required to ensure clinical relevance of this approach. It has been shown that melanoma in particular may feature certain shared epitopes across patients which are derived from incomplete splicing processes, which may render these cancers more susceptible to RI-derived neoantigens^38, 39^. Similar approaches across different histologies will provide further clarity on the role and contribution of RI neoantigens to tumor immunity across cancer contexts. Currently, the availability of clinically annotated cohorts with high quality RNA sequencing limits the widespread application of this method. Future efforts to promote RNA sequencing of matched normal samples alongside tumors, as well as optimization of methods specifically for formalin-fixed, paraffin-embedded tissue samples, will provide further clarity on the importance of specific intron retention events and corresponding neoantigens. Prediction of patient-specific RI-neoantigens has the potential to contribute to the development and further improvement of personalized cancer vaccines.

## Methods

### Clinical cohorts

Analysis was conducted on published cohorts of melanoma patients treated with immune checkpoint inhibitors. The Hugo et al. cohort included samples from 27 melanoma patients (26 pretreatment, 1 on-treatment) treated with the PD-1 inhibitor pembrolizumab^6^. Patient outcomes were classified as responding to therapy (R) (n=14) or not responding to therapy (NR) (n=13), as described in the original publication. These samples were sequenced from fresh frozen tissue using a standard, poly(A) selected protocol (personal communication, Willy Hugo). The Snyder cohort included post-treatment samples for 21 melanoma patients treated with ipilimumab (anti-CTLA-4 therapy)^5, 40^. Outcomes were classified as receiving long-term clinical benefit (LB) (n=8) or not receiving clinical benefit (NB) (n=13), as described in the original publication. RNA sequencing of the Snyder cohort was performed on fresh frozen tissue using a standard, poly(A) selected protocol.

### RI neoantigen pipeline

Raw RNA-Seq FASTQ files were pseudoaligned to an augmented hg19 (GENCODE Release 19, GRCh37.p13)^41^ transcriptome index containing both exonic and intronic transcript sequences, and transcript expression was quantified via kallisto^21^. The KMA algorithm^22^, implemented as a suite of Python scripts within an R package, was used to identify the genomic loci of expressed intron retention events with limited false positives. Using these RI loci, the UCSC Table Browser database^23^ was queried via public MySQL server to obtain the nucleotide sequences corresponding to the intronic regions and fragments of the previous exonic sequences, as well as the open reading frame orientation at the start of the intron. RI peptide sequences of 9-10 amino acids, with at least one intronic amino acid, were generated by translating open reading frames into intronic sequences until hitting an in-frame stop codon. These peptides, along with sample HLA Class I alleles identified via the POLYSOLVER algorithm^26^, were assessed for putative peptide-MHC I binding affinity via NetMHCPan v3.1^24^. A threshold of rank < 0.5% was used to identify putative RI neoantigens.

Several filters were applied at various steps throughout the pipeline to eliminate likely false positive RIs and RI neoantigens. After expression quantification, RIs expressed at a level ≤ 1 transcript per million, likely artifactual, were eliminated from the analysis. Additional expression-based filters were applied within the KMA algorithm: RIs that did not reach a level of at least five unique counts and whose neighboring exons did not reach a level of at least one transcript per million in at least 25% of samples in a cohort were eliminated as false positives. Due to the absence of matched normal RNA-Seq data for our melanoma clinical cohorts, a ‘panel of normals’ approach was taken in an attempt to filter out introns commonly retained in normal skin tissue, which would not produce immunogenic peptides due to likely host immune tolerance. RIs were identified in six normal skin samples (three individuals, two samples per individual: Individual ERS326932 with samples ERR315339 and ERR315376, Individual ERS326943 with samples ERR315372 and ERR315460, and Individual ERS327007 with samples ERR315401 and ERR315464) from the Human Protein Atlas^42^. RNA-Seq paired-end FASTQ files for each sample were downloaded from the following open-access link: https://www.ebi.ac.uk/arrayexpress/experiments/E-MTAB-1733/samples/. All normal sample retention profiles were highly concordant, both within and across individuals (Supplementary Fig. S6A). The final filter set of 7,050 normal RIs was obtained by intersecting the sets of RIs shared by each unique combination of one sample per individual—eight groups total (Supplementary Fig. S6B, Supplementary Table S7). These RIs were eliminated from downstream tumor sample analyses. In addition, RI peptides with amino acid sequences present in the normal proteome, derived from the UniProt human reference proteome version 2017_03, downloaded on 07/05/2017^25^, were filtered due to likely host immune tolerance. Finally, a set of RIs that were flagged due to abnormally high expression values and discovered upon manual review via Integrative Genomics Viewer^43^ to be erroneously-annotated in either the reference transcriptome or the Table Browser database were eliminated from the analysis (Supplementary Fig. S7A-D, Supplementary Table S7).

Pipeline code is publicly accessible on GitHub at https://github.com/vanallenlab/retained-intron-neoantigen-pipeline.

### Clinical cohort somatic neoantigen analysis

Putative somatic neoantigens were identified *in silico* for each sample as described in Van Allen et al. 2015. Briefly, BAM files from each cohort underwent sequencing quality control to ensure concordance between tumor and matched normal sequences and adequate depth of sequencing coverage. Single nucleotide variants were called using MuTect^44^ and insertions and deletions were called using Strelka^45^. Annotation of identified variants was done using Oncotator (http://www.broadinstitute.org/cancer/cga/oncotator). Sequences of 9-10 amino acid peptides with at least one mutant amino acid were generated. These peptides, along with HLA Class I alleles called with POLYSOLVER^26^ were analyzed using NetMHCpan v3.0^24^ to identify HLA-peptide binding interactions. For each patient, all peptides with predicted binding rank ≤ 2.0% for at least one patient HLA Class I allele were called somatic neoantigens.

### Responder-exclusive retained intron neoantigens

Responder-exclusive RI neoantigens were defined as neoantigens that were present in at least one patient who responded to immune checkpoint blockade therapy and absent in all nonresponders, using clinical annotations as published for both cohorts. RI transcripts yielding response-associated RI neoantigens were often expressed in multiple patients— both responders and non-responders— who lacked HLA alleles with predicted high binding affinity to the RI peptide. However, RI neoantigens were only classified as responder-exclusive if they were not expressed in any non-responders expressing HLA alleles with predicted high RI neoantigen binding affinity.

### Cell line analyses

Raw RNA-Seq data from the following published^8^ cell lines: CA-46, DOHH-2, HL-60, THP-1, MeWo, SK-Mel-5 were obtained from the Cancer Cell Line Encyclopedia^7^ via the NCI Genomic Data Commons^46^ and run through our computational pipeline as previously described, with minor adaptations as described henceforth. HLA Class I alleles were used for each cell line as enumerated in publication. A threshold of predicted binding rank ≤ 2.0% for at least one HLA Class I allele was used to distinguish cell line RI neoantigens. All pipeline filters applied to patient data described above were implemented on the cell line data *except* RI neoantigens expected to be retained in normal tissue were not filtered due to the fact that these experiments were focused on presentation of RI neoantigens rather than immune system stimulation once presented.

Mass spectrometric data from Ritz et al.^8^ as well as previously unpublished data for cell lines MeWo, DOHH2, and SKMEL5 was searched against a database consisting of 93,250 sequences of the human reference proteome downloaded from UniProt on July 7, 2017 concatenated with putative retained intron sequences (TPM > 1), or concatenated with 133,811 intron sequences with TPM < 1 (not retained) as negative control. Fragment mass spectra were searched with SEQUEST and filtered to a 1% false discovery rate with percolator to identify high confidence events.

### Gene set enrichment analysis

Gene expression was quantified in patient samples using kallisto^21^. Gene set enrichment analysis (GSEA)^30^ was run to compare both top quartile vs. bottom quartile RI load patients and immunotherapy responders vs. nonresponders. Initially, 50 Hallmark gene sets were tested. GSEA analyses of the Founders gene sets underlying the Hallmark gene sets that were significantly enriched in both top quartile vs. bottom quartile RI load patients and immunotherapy responders vs. nonresponders were subsequently performed. All statistical values reported are Benjamini-Hochberg FDR q values corrected for multiple hypothesis testing.

### Statistical analyses

Assessment of difference in means or medians for a continuous variable between two clinical response groups (i.e., clinical benefit vs. no clinical benefit) was performed using the nonparametric Mann-Whitney U test for non-normally-distributed variables (e.g., RI neoantigen burden). In the case of responder-exclusive RI neoantigen analyses, empirical p-values representing the percentage of time the outcome was observed in the setting of random phenotypic assignment were derived from 10,000 simulations with random permutation of the clinical benefit vs. no clinical benefit labels. To be specific, in each simulation round, the number of responder-exclusive RI neoantigens present in X samples, where 1 ≤ X ≤ total number of responders, was calculated. Then, empirical one-sided p-values were calculated as the proportion of simulations in which the outcome in question, or any greater outcome, was observed. All statistical analyses were conducted in the R statistical software environment (v.3.3.1).

## Data availability

Raw RNA-Seq data for the Snyder et al. 2014 patient cohort are available on dbGaP under accession code phs001038.v1.p1 and for the Hugo et al. 2016 cohort on the Sequence Read Archive (https://www.ncbi.nlm.nih.gov/sra) under the accession number SRA: SRP070710.

### Authors’ Contributions

**Conception and design**: A. C. Smart, C. A. Margolis, E. M. Van Allen Development of methodology: C. A. Margolis, A. C. Smart, H. Pimentel, M. X. He, T. Fugmann, D. Miao, K. Wong, E. M. Van Allen

**Analysis and interpretation of data (e.g., pipeline development, statistical analysis, computational analysis)**: C. A. Margolis, A. C. Smart, D. Adeegbe

**Writing, review, and/or revision of the manuscript**: C. A. Margolis, A. C. Smart, H. Pimentel, M. X. He, D. Miao, D. Adeegbe, T. Fugmann, K. Wong, E. M. Van Allen

**Study supervision**: E. M. Van Allen

## Acknowledgements

We are grateful to Dario Neri for fruitful discussions, Danilo Ritz for the purification of HLA peptides from cell lines, and Mahmoud Ghandi for assistance in coordinating access to cell line transcriptome data.

## Supplementary Materials

Figures S1-S7

Tables S1-S7

